# Interpreting prediction intervals and distributions for decoding biological generality in meta-analyses

**DOI:** 10.1101/2024.09.10.612386

**Authors:** Yefeng Yang, Daniel W. A. Noble, Alistair M. Senior, Malgorzata Lagisz, Shinichi Nakagawa

## Abstract

Despite the importance of identifying predictable regularities for knowledge transfer across contexts, the generality of ecological and evolutionary findings is yet to be systematically quantified. We present the first large-scale evaluation of generality using new metrics. By focusing on biologically relevant study levels, we show that generalization is not uncommon. Overall, 20% of meta-analyses will produce a non-zero effect 95% of the time in future replication studies with a 70% probability of observing meaningful effects in study-level contexts. We argue that the misconception that generalization is exceedingly rare is due to conflating within-study and between-study variances in ecological and evolutionary meta-analyses, which results from focusing too much on total heterogeneity (the sum of within-study and between-study variances). We encourage using our proposed approach to elucidate general patterns underpinning ecological and evolutionary phenomena.

---

Ecologists and evolutionary biologists strive to uncover predictable regularities about the biological world, seeking biological generality ^1^. Unveiling general patterns that underpin a given phenomenon is of immense interest in ecology and evolution ^2^. This pursuit enables stakeholders such as practitioners and policymakers to transfer knowledge across diverse populations and contexts, including different ecosystems, species, and spatio-temporal scales. This, in turn, enhances predictive capabilities and facilitates more precise management, intervention, and conservation practices. Meta-analyses play a pivotal role in this process. By synthesizing evidence from a collection of comparable study systems ^3^, meta-analyses can scrutinize the extent to which inferences drawn from a specific population can be replicated (replicability), extended beyond the reference population to a new population of interest (transferability), and extrapolated to a broader target population (generalizability) as requested by stakeholders ^1,4^.

Despite the high heterogeneity revealed in ecological and evolutionary findings ^5^, the generality of these findings has yet to be empirically quantified, despite its critical importance. The ‘generality gap’ likely stems from a lack of suitable metrics for quantifying generality. Traditionally, heterogeneity—reflecting variability in population (true) effect sizes— has been used as a proxy for inferring the generality of effects in a population of studies ^6^. Zero heterogeneity indicates population effect sizes are consistent across different studies, implying that the population effect is perfectly generalizable, transferable, and replicable across different contexts. While heterogeneity metrics, such as Cochran’s *Q* and *I*^2^, are commonly used for this purpose, it is challenging to interpret and understand them as indicators of generality^6^. More importantly, the current practice of estimating total heterogeneity to infer overall generality ^7^ completely ignores the hierarchical nature of ecological and evolutionary data structures ^8,9^, leading to the illusion that little to no generality can be drawn from ecological and evolutionary findings ^5^. Explicit partitioning of heterogeneity at different hierarchical levels (e.g., study- and species-levels) provides novel insights for understanding the generality of population effects tailored to specific population characteristics such as taxa, ecosystems, locations, and times ^7^.

Here, we present the first systematic investigation to demonstrate how the use of total heterogeneity, which confounds different sources of heterogeneity, has impeded our ability to generalise ecological and evolutionary findings. We show how focusing on the decomposition of heterogeneity can provide more appropriate predictions of generalisability. To achieve this, we use a dataset comprising 512 meta-analyses with 109,495 observed effect sizes ^10,11^, and we apply hierarchical partitioning techniques (see below and Methods) to estimate prediction intervals (PIs) and derive predictive distributions (PDs). PIs and PDs both offer direct measures of generality ^12,13^. This is because PIs provide an interval that indicates the extent to which the phenomenon of interest (more precisely, the population effect) can be generalized in replication studies with a certain probability (usually 95%) ^12^. PDs offer a probabilistic estimate of the entire distribution of effect sizes from new studies ^13^, enabling the estimation of the likelihood of observing future studies from similar contexts above a biologically or practically meaningful threshold. For example, consider a situation where a conservation biologist is interested in knowing how frequently a conservation intervention in a replicated study would surpass an SMD = 0.5. If the probability of observing an effect at the between-study level is 88% then we can conclude that 88% of future studies are likely to produce an SMD value of 0.5 or greater (Fig. 1), with strong implications for management, conservation practices, and policymaking.

**Fig. 1.**
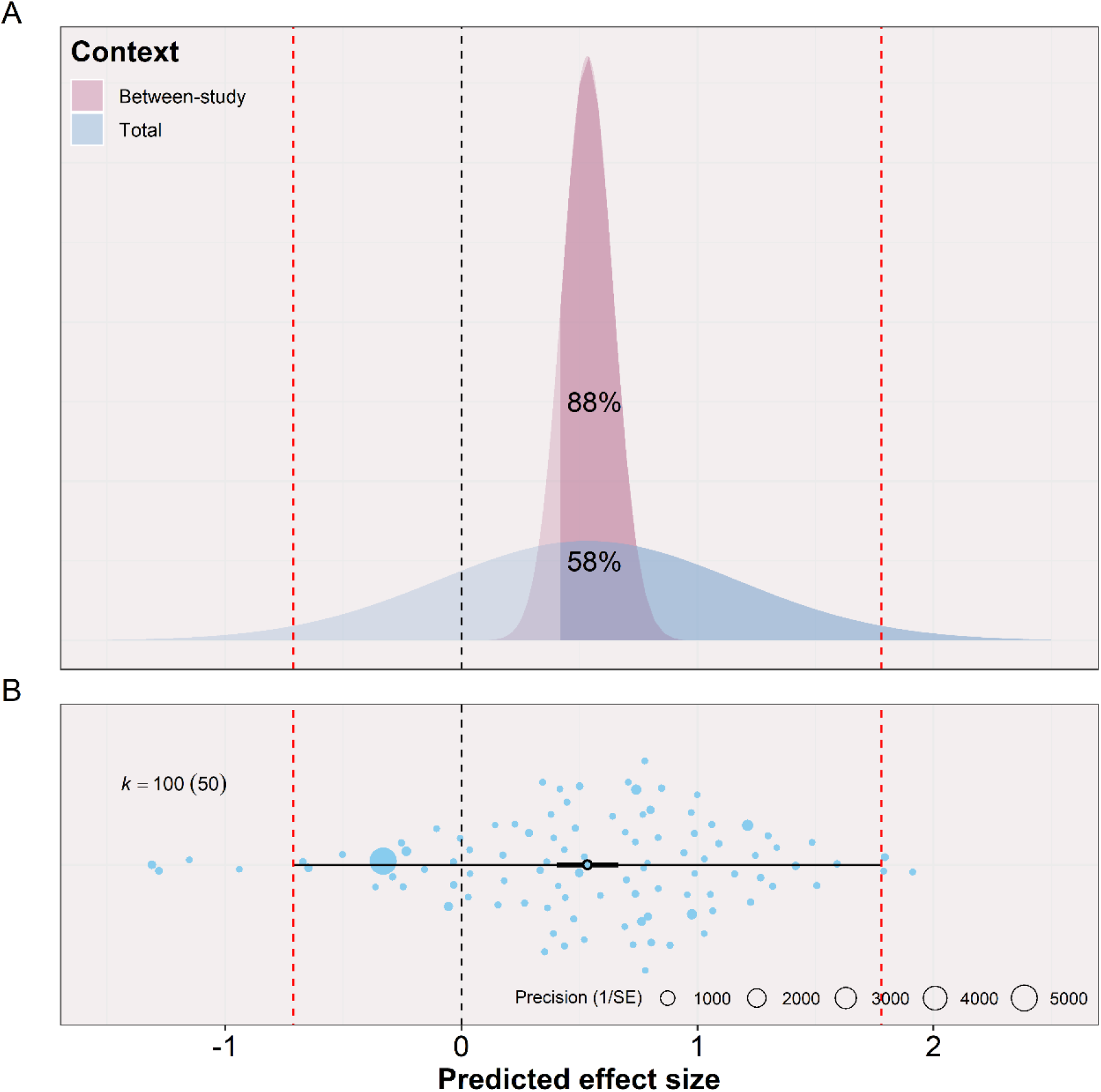
Example of how prediction distributions (PDs; panel A) combined with prediction intervals (PIs; panel B) can be used to understand the generality of population effect size. We simulated hierarchical data with three sources of variance [between-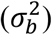 and within-study 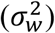 variance as well as sampling error]. *k* = 100 effect size estimates were nested within a total of 50 studies, with a 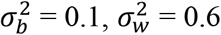, and an overall mean population effect µ_*θ*_ of 0.5. The top plot (Panel A) provides the probability of observing an effect from a new study above the meaningful threshold (i.e., the lower boundary of 95% CIs around the mean). Red dashed lines are the upper and lower 95% PI, and black dashed lines are the ‘null’ effect. The bottom plot (Panel B) displays the estimated overall mean population effects, 95% confidence intervals (CIs), and 95% PI. The PDs were constructed using a *t-*distribution with *k* – 1 degrees of freedom, µ_*θ*_ as the location parameter, and total 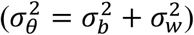 variance or study-specific variance 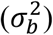 as the scale parameter.

We assessed the ‘generalisability’ of a given meta-analysis by deriving PIs and PDs using 1) total heterogeneity and 2) after partitioning heterogeneity to within- and between study levels. We then compare how conclusions about generalisability change when comparing these scenarios. The procedure of “decomposition” is logical and straightforward. Suppose *θ*_*ij*_ is the *i*-th population effect size in *j*-th study included in a meta-analysis. We first fit the three-level meta-analytic model ^14,15^ to each of the 512 meta-analyses to estimate the average effect size in the population µ_*θ*_ = *E*[*θ*_*ij*_] (i.e., meta-analytic overall mean), and variance 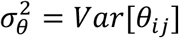 of the population effect size *θ*_*ij*_. A typical assumption of the three-level meta-analysis model implies that *θ*_*ij*_ is a random sample from a distribution of population effect sizes with mean µ_*θ*_ and variance 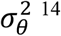. Based on the estimated parameters from these models, we employed both integration methods and Monte Carlo simulations (as detailed in **Methods**) to derive PIs and PDs from the distribution of population effect sizes for each meta-analysis. The three-level model has two properties that provide the statistical foundation for us to test the generality of meta-analysis results: (1) inferences and predictions can be made for studies that are not included in the meta-analysis, (2) variance 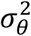 can be decomposed into between- and within-study level variances (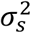 and 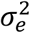), enabling us to disentangle the generality at specific contexts.

Overall, 321 out of 512 (63%) meta-analyses showed a statistically significant average population effect (non-zero effect; Table S1), where the null hypothesis µ_*θ*_ = 0 was rejected at a significance level of 0.05, or equivalently, the 95% confidence interval (CI) excluded the null effect (zero). This finding indicates that for every negative (null) ecological and evolutionary meta-analytic finding, there are approximately two positive ones. However, µ_*θ*_, the uncertainty in µ_*θ*_ (95% CI), and the hypothesis test (p-value) are insufficient to summarize the generalisation of the heterogeneous effects ^1^. To better assess generalisability, we should construct and report on PIs. PIs derived using total heterogeneity 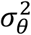 and sampling errors (see **Methods**) revealed that only 21 excluded the null effect within their 95% PIs among the 321 meta-analyses showing statistically significant effects (Table S1). Therefore, only a very small number of meta-analyses offer assurance that future replication studies (studies on similar topics) would yield a non-zero effect.

However, this seemingly low degree of generalizability partially results from not correctly accounting for the hierarchical nature of ecological and evolutionary data. Decomposing PIs at the between-study suggests 71 meta-analyses achieve generalisation at the between-study-level context (after controlling for the within-study variance; Table S1); in other words, among studies with statistically significant average population effects, 22% of future replication studies would produce a non-zero effect (95% of the time; Fig. 2A). Different effect size measures showed similar patterns (Table S2). Given these results, the use of total heterogeneity for PIs, which overlooks the hierarchical structure of data, creates the illusion that generalization is rare in ecological and evolutionary studies. However, by decomposing overall generalization at the between-study level—after accounting for within-study variability (see below for the approach)—we derived more biologically meaningful heterogeneity, which can be interpreted as generality among studies (populations). This finding implies that generality is more achievable among studies, including those in ecological and evolutionary meta-analyses than previously thought.

**Fig. 2.**
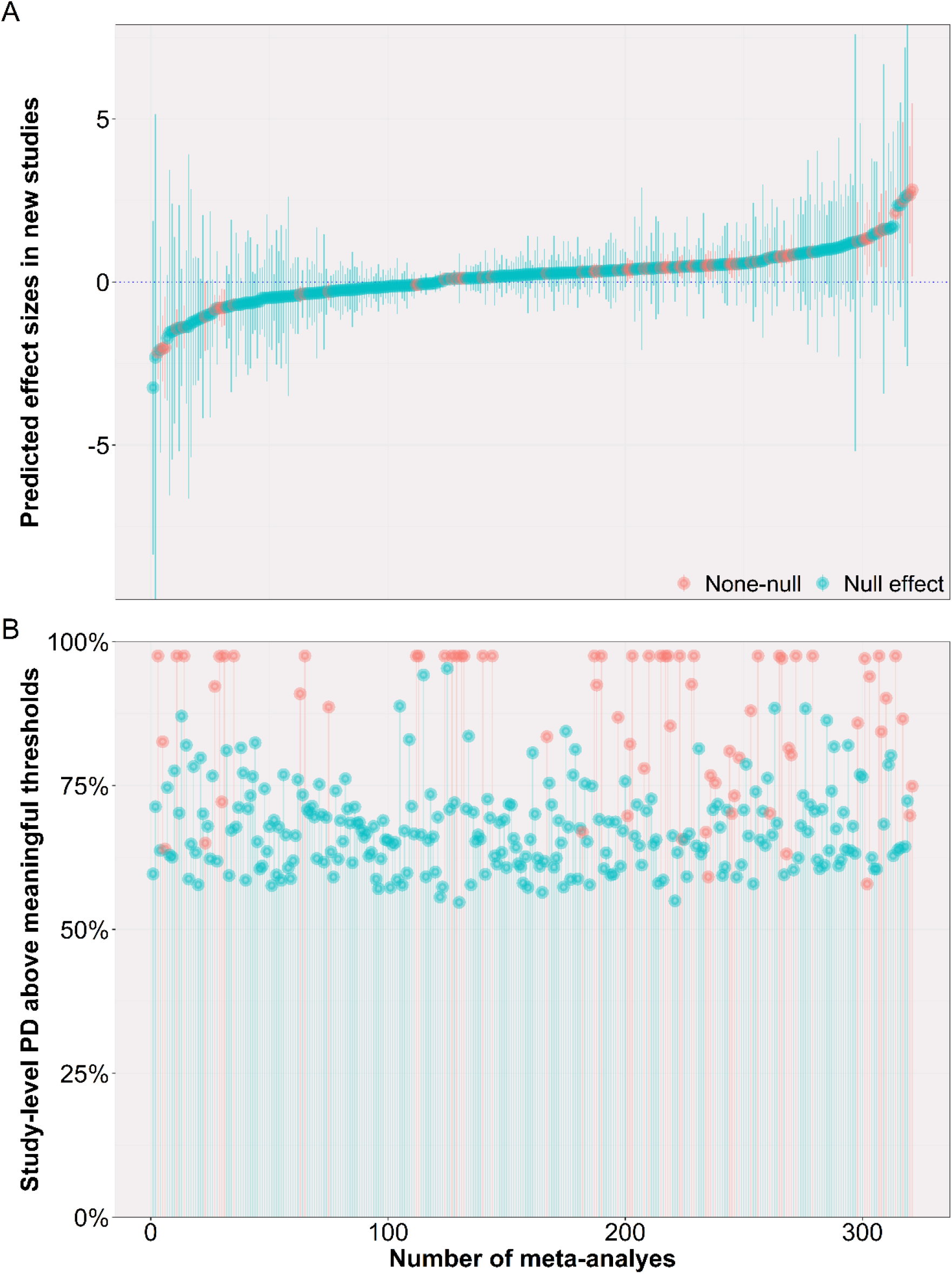
The overall and study-specific generality of 247 meta-analyses with statistically significant overall mean effects. The generality is measured as 95% prediction intervals (PIs; Panel A) and the probability of observing an effect from a new study above a practically meaningful threshold (Panel B) at the study level. 95% of effect sizes from future studies from similar contexts will yield statistically significant effects if PIs exclude the null effect. Prediction intervals (PDs) offer the estimation of the likelihood that the effect exceeds a biologically or practically meaningful threshold (in this case, the lower confidence limit; see the main text for the sensitivity analysis). Each dot in Panel A represents each meta-analysis’s average population effect size. The whisker denotes 95% PIs. Each dot in Panel B represents the probability of future studies from similar contexts exceeding the lower bound of 95% confidence intervals. Note that practically meaningful thresholds can be adjusted according to practitioners’ needs (e.g., the smallest effect size that can trigger importance differences ^17,18^).

This finding is particularly striking given that ecological and evolutionary meta-analyses often synthesize evidence from highly heterogeneous contexts, such as large spatial scales, different time periods, multiple systems, and a diverse array of organisms ^3^. Importantly, the emerging “big-team” open science approach in ecology and evolutionary biology offers effective ways to control for within-study variance. For example, big-team science initiatives such as the ManyBirds project (https://themanybirds.com/) and coordinated distributed experiments ^16^ often employ standardized, controlled protocols to advance our understanding of general principles in ecology and evolution.

Statisticians have emphasised the importance of computing the probability density to accurately capture the likelihood of each effect size within the intervals ^19,20^. By considering the entire distribution of population effects, PDs offer more holistic information for measuring generality. We found that the probability of observing an effect from a new replication study above a meaningful threshold, defined as the lower boundary of 95% CIs (see below for the sensitivity analysis), was 64% (Fig. 2, Fig. S1, and Table S3). Different types of effect sizes, such as standardized mean difference (e.g., Cohen’s *d*), log response ratio, and Fisher’s *Zr*, showed similar patterns (Fig. S2 – S4, and Table S4). The study-level probability of observing effects above meaningful thresholds increased to 71% after controlling for within-study-level variances (Fig. 2, and Fig. S1). Therefore, the results of PDs corroborated those from PIs; while the overall generality is rare, generality at the study-level context is not uncommon in ecological and evolutionary meta-analyses. A potential limitation of the above findings is that PIs are not probability intervals, giving the impression that all effect sizes within the upper and lower intervals are equally likely ^21^.

While we used the lower confidence limit of a meta-analysis as a general proxy for a meaningful threshold, the smallest effect size of interest (SESOI) could serve as a more appropriate lower bound for meaningful effects. However, determining SESOI requires expert knowledge and is likely topic-dependant (see Methods) ^18^. For example, in medical research, methods have been proposed to establish the smallest effect that patients perceive as beneficial, also known as the minimally clinically important difference ^17^. Therefore, it is recommended to use the SESOI specific to the phenomenon of interest when computing PDs once such SESOI is well-developed and understood for a certain topic. As a sensitivity analysis, we used conventional cutoffs for the ‘small effect’ (0.2 for Cohen’s *d*, and 0.1 for Fisher’s *Zr*) ^22^ as the SESOI. We found that the study-level probability of observing effects above meaningful thresholds was 51% for Cohen’s *d* and 64% for Fisher’s *Zr* (Fig S5 and Table S5).

Conventional wisdom assumes that highly heterogeneous population (meta-analytic overall) effects are contextually sensitive, automatically yielding non-generalizable results ^1^. However, our study reveals that mainstream heterogeneity practices ^7^, which rely on total heterogeneity, often wrongly conclude that meta-analytic effects are non-generalisable. The heterogeneity relevant to the generalization of population effects should be defined at biologically meaningful levels, such as the study level (population), rather than at the total level, which mixes up within-study-level heterogeneity, which could be standardised by, for example, employing unified protocols (e.g., big-team science). The “decomposition” technique we present here allows us to partition total heterogeneity into study-level heterogeneity, thus examining the generalization at biologically meaningful levels. Our results indicate that achieving generality at the study level is feasible and that the generalisability of meta-analytic findings is likely underestimated. This approach can also be extended to estimate generalization at meaningful levels beyond the study level. For example, total heterogeneity can be partitioned further into species or geographic location levels, providing us with the chance to quantify the degree of generalization at these two levels beyond those we showed here ^23^. To facilitate the exploration of generalisation, we provide all R scripts with custom functions. These tools will help ecologists and evolutionary biologists assess and interpret the generality of their meta-analytic findings more accurately, ultimately contributing to uncovering predictable regularities that enable practitioners and policymakers to generalize and transfer knowledge across populations and contexts.

## Methods

### Datasets

The ecological and evolutionary database (522 meta-analysis datasets) used in this study was initially compiled by Costello ^11^, O’Dea ^10^, and their colleagues. They systematically searched for meta-analysis papers published in ecological journals, including those from the Ecological Society of America and journals of the British Ecological Society. Additionally, they supplemented the database with high-profile journals, such as Nature, and Science. We excluded datasets where the multilevel model did not achieve convergence. Despite adjusting key parameters for optimizing the log-likelihood function ten datasets failed to converge (details provided below). As a result, our database comprised 512 meta-analysis datasets, which included 17,770 primary studies and 109,495 effect size estimates. Various effect size measures were included in our database, including standardised mean difference (*N* = 223), log response ratio (*N* = 128), Fisher’s *Zr* (*N* = 128), and others (e.g., odds ratio, regression coefficient). On average, each meta-analysis dataset contained 240 effect size estimates from 40 studies, with median values of 64 and 23, respectively.

We fit a multilevel meta-analytic model with three levels, with the effect size estimate *ES*_[*i*]_ being expressed as a combination of the population mean effect (µ_*θ*_; overall mean effect size), along with random effects at two levels (between- and within-study), and sampling error:

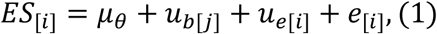

The typical assumptions for this equation are: (i) the between-study random effect *u*_*b*[*j*]_ is assumed to follow a normal distribution with a mean of zero and variance 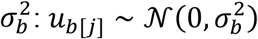, (ii) the within-study random effect *u*_*e*[*i*]_ is assumed to follow a normal distribution with a mean of zero and variance 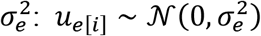, and (iii) sampling error *e*_[*i*]_ is assumed to follow a normal distribution with a mean of zero and a variance defined by the sampling variance (𝒱 _[*i*]_) corresponding to each effect size, *i*, such that *e*_[*i*]_ ∼ 𝒩 (0, 𝒱_[*i*]_). The assumption of homogeneous variances for the random effects can be relaxed to account for heteroscedasticity ^24^. Similarly, the assumption of independent sampling errors (*e*_[*i*]_) can be relaxed to allow for sampling error covariance 𝒱_[*i*]_ ^14^. Our multilevel meta-analytic model decomposes the total variance 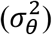 around the true effect into between-study level variance 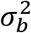 and with-study level variance 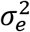, allowing us to examine the generality among studies (populations). Note that the multilevel meta-analytic model can be extended to incorporate the random effects that have biological interest. For example, the multilevel meta-analytic model can include species or geographic location as the additional random effects, providing us with the chance to quantify the degree of generalization at these two levels.

We employed the *rma*.*mv()* function from the *metafor* package ^15^ to fit all 512 meta-analysis datasets to the three-level meta-analytic model (Equation 1). We used restricted maximum likelihood (REML) as the variance estimator and the quasi-Newton method to optimize the likelihood function over variance estimation (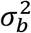 and 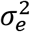), with a threshold of 10^−8^, a step length of 1, and a maximum iteration limit of 1,000. All models successfully converged under these settings. We confirmed the identifiability of variance estimation (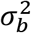 and 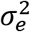) by checking their likelihood profiles (i.e., the peak of the curve showing the restricted log-likelihood as a function of variance 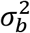 and 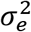).

### Prediction intervals

Prediction intervals (PIs) are underreported but insightful in understanding meta-analytic heterogeneity and generality. For example, surveys have shown that less than one percent (1/102) of studies report PIs ^25^. PIs are derived from 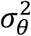 and provide a range within which a future effect size is predicted to fall with a certain probability ^12^, often 95%. For example, consider a conservation intervention with a mean effect size (SMD) of -0.5 and 95% PIs of [-0.2 to -0.8]. This indicates that 95% of future interventions implemented are predicted to decrease the conservation outcomes of interest by between 0.2 to 0.8 standard deviations. Unlike the point estimate of heterogeneity, such as total variance 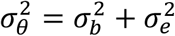, PIs offer an interval to inform the extent to which the focal effect can be generalized ^26^. 95% of effect sizes from future studies from similar contexts will yield statistically significant effects if PIs exclude the null effect. Under Equation 1, 95% PIs at the total level can be computed by ^14^:

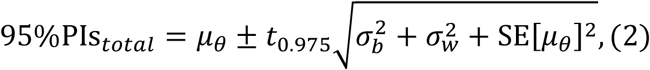

where *t*_0.975_ denotes the 97.5*th* percentile of a *t*-distribution (with *k*−1 degrees of freedom ^27^, where *k* is the number of sample size), and SE[µ] denotes the standard error of the mean effect µ_*θ*_. Study level 95% PIs can be obtained by controlling for the within-study variance 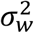:

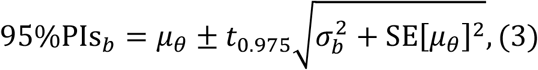

### Prediction distributions

We further propose using predictive distribution (PDs) to show the entire distribution of true effects while accounting for statistical noise, offering a more holistic measure of generality. In the Bayesian framework, PDs, known as posterior distributions, are a natural part of the analytical approach. However, even frequentist approaches can adopt PDs (sometimes referred to as “empirical Bayes”) to achieve similar aims ^28^. An advantage of the PD is its ability to calculate the probability that a true effect size exceeds a biologically or practically meaningful threshold. For a given threshold of practically meaningful effect size (*q*), the estimated proportion of true effect sizes above this threshold has been proposed as a measure of heterogeneity ^13^. In our study, we extended this approach to quantify the probability of observing an effect from a new study above a practically meaningful threshold (*q*), *P*(*x* > *q*). By definition, *P*(*x* > *q*) serves as a direct measure of the generalizability of an effect of interest. Consider a case that 69% of effect sizes representing the efficacy of a conservation intervention are predicted to surpass a threshold value representing a practically significant effect (Fig. 1, where we assumed the lower confidence limit representing the threshold; see below). Assuming similar configurations of study contexts in the sampled future cases, we can infer that the intervention will achieve this benefit in 69% of future cases, with strong implications for policymaking. To compute the probability the probability *P*(*x* > *q*) of the true effect in a new study will exceed the meaningful threshold *q*, we employed two approaches, which were detailed below along with a discussion on how to define a practically meaningful threshold *q*.

### Monte Carlo simulations of future effect sizes specific to each meta-analytic context

Equations 2 and 3 imply that effect sizes from new studies follow a *t*-distribution with *k*−1 degrees of freedom ^27^ (where *k* denotes the number of sample size). Based on REML estimates of model parameters (Equation 1), we recovered the corresponding *t*-distribution for each meta-analytic context and then sampled 10 ^5^ effect sizes from this distribution, specific to each context (in terms of statistical properties). By definition, these 10 ^5^ effect sizes represent potential outcomes of future studies that maintain similar design configurations as those covered in the given meta-analysis. To sample the 10 ^5^ effect sizes from the *t*-distribution with *df* = *k* − 1 degrees of freedom, we used the following formula ^29,30^:

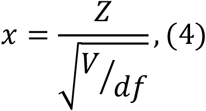

where *Z* is a random variable drawn from a standard normal distribution *N*(0,1), and *V* is a random variable drawn from a chi-squared distribution with *df* degrees of freedom χ^2^(*df*). The random variable *x* (Equation 4) has the probability density given by:

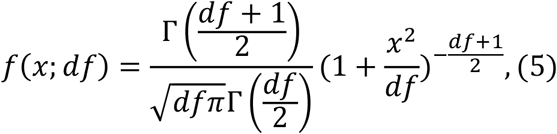

where *x* is the value at which the density (height of the probability density function) is evaluated, and Γ is the gamma function. After sample *x* = 10 ^5^ effect sizes for each meta-analysis, we estimated *P*(*x* > *q*) by calculating the proportion of sampled effect sizes that exceed the practically meaningful threshold *q*.

### Integration over the probability density function established from the estimated model parameters

Alternatively, we estimated *P*(*x* > *q*) by integrating over the probability density function derived from the estimated model parameters. To facilitate the computation, we transformed the *t*-distribution with *df* = *k* − 1 degrees of freedom into a location-scale distribution characterized by location parameter µ_*θ*_ and scale parameter σ_*θ*_. These parameters µ_*θ*_ and σ_*θ*_ of each meta-analysis were obtained via the REML estimator in Equation 1. The location-scale transformation of the *t*-distribution with *df* = *k* − 1 degrees of freedom is given by:

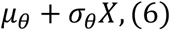

Then, the probability density function of the location-scale *t*-distribution is expressed as ^31^:

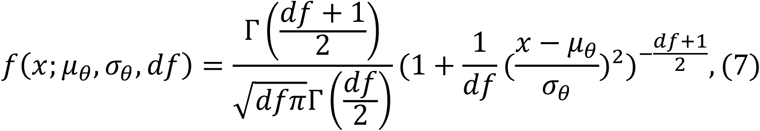

For each meta-analysis, after determining µ_*θ*_, σ_*θ*_, and *df*, we integrated *f*(*x*; µ_*θ*_, σ_*θ*_, *df*) from Equation 7 to calculate *P*(*x* > *q*). Essentially, *P*(*x* > *q*) corresponds to the area under the upper tail of the cumulative distribution function, which can be computed as 1 − *P*(*x* < *q*), where *P*(*x* < *q*) represents the lower tail area. Leveraging the decomposition technique outlined earlier, we also computed study-level *P*(*x* > *q*) based on the *t*-distribution location parameter µ_*θ*_ and scale parameter σ_*b*_.

### Practically meaningful effect size thresholds

Determining the specific, practically meaningful threshold (*q*) for each of the meta-analyses in our study is challenging, requiring domain-specific knowledge, insights from practitioners, and relevant considerations of practical relevance. In fields such as medicine and psychology, researchers have proposed using the smallest effect size of interest (SESOI) as the lower bound for a meaningful effect that (i) can trigger important differences or is practically and theoretically meaningful ^17,18^. Several approaches can help establish SESOI, such as anchor-based methods or distribution-based methods ^18^. However, a consensus has yet to be reached regarding the most suitable methodology. For example, anchor-based methods are limited by the choice of an anchor question, while distribution-based methods rely purely on statistical reasoning. Moreover, it is challenging, if not impossible, to use these methods to establish SESOI for each of the meta-analyses because no well-performed methods have been extended to the field of ecology and evolution, and our database needed more study context information to support the computation. In contrast, we used the lower confidence limit of each meta-analysis as a general proxy for the meaningful threshold. The use of the SESOI specific to each meta-analysis is recommended when computing PDs once approaches for establishing the SESOI are well-developed in fields like ecology and evolution. Furthermore, we provide all reproducible R scripts, which makes updating the computation of PDs based on specific SESOI feasible once it becomes available.

## Data and code availability statement

The data and code to reproduce the results of this study are available in the GitHub repository (https://github.com/Yefeng0920/generality_PD). We will archive the data and code to a permanent repository (e.g., Zenodo or Figshare) after acceptance.

## Acknowledgments

Thanks to Rebecca Spake for her previous comments and suggestions on this manuscript. YY was funded by the National Natural Science Foundation of China (NO. 32102597). ML and SN were funded by the Australian Research Council Discovery Grant (DP210100812 & DP230101248). SN was also supported by a Canada Excellence Research Chair (CERC-2022-00074). AMS is funded by an Australian Research Council Future Fellowship (FT230100240).

## Author contributions

All authors were involved in the conceptualisation of the study and editing of the manuscript. Yefeng Yang collected the data, analysed the data, and drafted the manuscript with the help of Shinichi Nakagawa. Alistair M. Senior suggested the sensitivity analysis. Daniel W. A. Noble assisted in the development of the R script for visualization.

## Additional information

Supplementary material: Extended Figures and Tables

Supplementary data

## Reference

1 Spake, R. et al. Improving quantitative synthesis to achieve generality in ecology. Nature Ecology & Evolution, 1-11 (2022).

2 Lawton, J. H. Are there general laws in ecology? Oikos, 177-192 (1999).

3 Gurevitch, J., Koricheva, J., Nakagawa, S. & Stewart, G. Meta-analysis and the science of research synthesis. Nature 555, 175–182 (2018).

4 Martin, P. A. et al. Flexible synthesis can deliver more tailored and timely evidence for research and policy. Proceedings of the National Academy of Sciences 120, e2221911120 (2023).

5 Senior, A. M. et al. Heterogeneity in ecological and evolutionary meta-analyses: its magnitude and implications. Ecology 97, 3293–3299 (2016).

6 Higgins, J. P. & Thompson, S. G. Quantifying heterogeneity in a meta-analysis. Statistics in medicine 21, 1539–1558 (2002).

7 Yang, Y. et al. Measuring biological generality in meta-analysis: a pluralistic approach to heterogeneity quantification and stratification. (2023).

8 Nakagawa, S., Yang, Y., Macartney, E. L., Spake, R. & Lagisz, M. Quantitative evidence synthesis: a practical guide on meta-analysis, meta-regression, and publication bias tests for environmental sciences. Environmental Evidence 12, 8, doi:10.1186/s13750-023-00301-6 (2023).

9 Noble, D. W. et al. Meta-analytic approaches and effect sizes to account for ‘nuisance heterogeneity’in comparative physiology. Journal of Experimental Biology 225, jeb243225 (2022).

10 O’Dea, R. E. et al. Preferred reporting items for systematic reviews and meta-analyses in ecology and evolutionary biology: a PRISMA extension. Biological Reviews 96, 1695–1722 (2021).

11 Costello, L. & Fox, J. W. Decline effects are rare in ecology. Ecology 103, e3680 (2022).

12 IntHout, J., Ioannidis, J. P., Rovers, M. M. & Goeman, J. J. Plea for routinely presenting prediction intervals in meta-analysis. BMJ open 6, e010247 (2016).

13 Mathur, M. B. & VanderWeele, T. J. New metrics for meta-analyses of heterogeneous effects. Statistics in Medicine 38, 1336–1342 (2019).

14 Yang, Y., Macleod, M., Pan, J., Lagisz, M. & Nakagawa, S. Advanced Methods and Implementations for the Meta-analyses of Animal Models: Current Practices and Future Recommendations. Neuroscience & Biobehavioral Reviews, 105016 (2022).

15 Viechtbauer, W. Conducting meta-analyses in R with the metafor package. Journal of statistical software 36, 1–48 (2010).

16 Fraser, L. H. et al. Coordinated distributed experiments: an emerging tool for testing global hypotheses in ecology and environmental science. Frontiers in Ecology and the Environment 11, 147–155 (2013).

17 McGlothlin, A. E. & Lewis, R. J. Minimal clinically important difference: defining what really matters to patients. JAMA 312, 1342–1343 (2014).

18 Wang, Y. et al. A step-by-step approach for selecting an optimal minimal important difference. BMJ 381 (2023).

19 Jackson, C. H. Displaying uncertainty with shading. The American Statistician 62, 340–347 (2008).

20 Barrowman, N. J. & Myers, R. A. Raindrop plots: a new way to display collections of likelihoods and distributions. The American Statistician 57, 268–274 (2003).

21 Bishop, J. & Nakagawa, S. Quantifying crop pollinator dependence and its heterogeneity using multi-level meta-analysis. Journal of Applied Ecology 58, 1030–1042 (2021).

22 Cohen, J. Statistical power analysis for the behavioral sciences. 2nd edn, (Routledge, 1988).

23 Yang, Y. et al. Species sensitivities to artificial light at night: A phylogenetically controlled multilevel meta-analysis on melatonin suppression. Ecology Letters 27, e14387 (2024).

24 Viechtbauer, W. & López-López, J. A. Location-scale models for meta-analysis. Research synthesis methods 13, 697–715 (2022).

25 Nakagawa, S. et al. The orchard plot: cultivating a forest plot for use in ecology, evolution, and beyond. Research Synthesis Methods 12, 4–12 (2021).

26 van Aert, R. C., Schmid, C. H., Svensson, D. & Jackson, D. Study specific prediction intervals for random-effects meta-analysis: A tutorial: Prediction intervals in meta-analysis. Research synthesis methods 12, 429–447 (2021).

27 Knapp, G. & Hartung, J. Improved tests for a random effects meta-regression with a single covariate. Statistics in medicine 22, 2693–2710 (2003).

28 Raudenbush, S. W. & Bryk, A. S. Empirical Bayes meta-analysis. Journal of educational statistics 10, 75–98 (1985).

29 Ruppert, D. & Matteson, D. S. Statistics and data analysis for financial engineering. Vol. 13 (Springer, 2011).

30 Johnson, N. L., Kotz, S. & Balakrishnan, N. Continuous univariate distributions, volume 2. Vol. 289 (John wiley & sons, 1995).

31 Jackman, S. Bayesian analysis for the social sciences. (John Wiley & Sons, 2009).

